# Chemical composition and antioxidant properties of Thai propolis: Role of caffeic acid phenethyl ester (CAPE)

**DOI:** 10.64898/2026.01.10.698791

**Authors:** Thanyarat Chuesaard, Veeranan Chaimanee, Jantipa Jobsri, Pattraporn Pukklay

## Abstract

Propolis, a resinous substance produced by *Apis mellifera*, is rich in polyphenols and flavonoids and exhibits diverse biological activities, including antioxidant and antimicrobial effects. This study aimed to characterize the chemical composition of Thai propolis and evaluate its free radical scavenging activity. Ethanolic (EEP) and aqueous (WEP) extracts were prepared and analyzed. Gas chromatography–mass spectrometry (GC-MS) identified major constituents of EEP, including phenolic acids, flavonoids, and sugars. Antioxidant activities were assessed using DPPH and hydrogen peroxide scavenging assays. EEP exhibited significantly higher radical scavenging activity than WEP, with maximum DPPH inhibition of 87.46% at 5 mg/mL. CAPE effectively scavenged superoxide and hydroxyl radicals and protected plasmid DNA from oxidative damage. These results highlight CAPE as bioactive compound contributing to the antioxidant properties of Thai propolis. Thai propolis, particularly its ethanolic extract, represents a promising natural source of antioxidants for potential applications in nutraceuticals, functional foods, and oxidative stress-related disease prevention.

## Introduction

Propolis is a resinous substance produced by *Apis mellifera*, composed of waxes, salivary enzymes, and plant bud exudates. Honeybees use propolis for nest construction, sealing openings, covering carcasses, and its antimicrobial properties [1,2]. The chemical composition of propolis varies depending on geographical location and ecological factors, and it is rich in polyphenols and flavonoids [3]. The bioactive constituents include phenolic esters [4], p-coumaric acid, diterpenes, lignans, and flavonoids [5,6]. Among these, caffeic acid phenethyl ester (CAPE), a derivative of p-coumaric acid, cinnamic acid, and caffeic acid, has attracted significant attention due to its bioactivity [7]. CAPE exhibits both cis- and trans-structures and has been incorporated into PLGA nanoparticles for enhanced delivery [8]. Flavonoids and CAPE have demonstrated cytotoxic and antiproliferative effects in cancer cell lines, including PC-13, DU-145, and PC-3 [9,10].

The analysis of phenolic compounds and their derivatives is critical for understanding the biological activities of propolis. In vitro studies have shown that propolis exerts anti-inflammatory effects on dental pulp cells [11], and ethanolic extracts are commonly used to assess prooxidant and free radical scavenging activities. Etha nolic propolis extracts have also displayed antibacterial activity against *Klebsiella pneumoniae*, *Pseudomonas aeruginosa*, and *Escherichia coli* [12], as well as antifungal activity against *Candida albicans*, *Cryptococcus neoformans*, and *Saccharomyces cerevisiae* [13]. Extraction methods using ethanol gradients are widely employed to selectively dissolve bioactive compounds while minimizing the co-extraction of inert molecules. Optimized extraction protocols enhance the yield of active compounds, reduce solvent usage, and decrease processing time [14].

Reactive oxygen species (ROS), including superoxide anion and hydroxyl radical, are highly reactive molecules that can damage lipids, DNA, and proteins [15,16]. Hydrogen peroxide, in the presence of transition metals such as Fe²⁺ and Cu²⁺, generates ROS that induce cellular injury and death. Cells employ antioxidant enzymes such as catalase to neutralize these reactive species. Given these oxidative challenges, natural antioxidants such as those found in propolis are of interest for mitigating ROS-induced damage.

In this study, Thai propolis from *A. mellifera* was analyzed to identify its main chemical components using gas chromatography–mass spectrometry (GC-MS). The antioxidant activities of ethanolic (EEP) and aqueous (WEP) extracts were assessed, and the bioactivity of CAPE against free radicals was further evaluated to elucidate its potential protective effects.

## Materials and Methods

### Chemicals and reagents

The following chemicals were used in this study: ethanol (RCI LabScan, Thailand); 2,2-diphenyl-1-picrylhydrazyl (DPPH; Sigma-Aldrich, Germany); hydrogen peroxide (Merck, USA); xanthine (Sigma, China); ethylenediaminetetraacetic acid (EDTA; Ajax Finechem, USA); nitroblue tetrazolium (Sigma, USA); sodium carbonate (Kemaus, USA); xanthine oxidase (Fluka, USA); deoxyribose (Sigma-Aldrich, China); ferrous sulfate (Himedia, USA); trichloroacetic acid and 2-thiobarbituric acid (Merck, Germany); plasmid DNA pBR322 (New England Biolabs, USA); copper(II) chloride (Kemaus, USA); L-ascorbic acid (Kemaus, USA); and quercetin (Sigma-Aldrich, USA). All reagents were of analytical grade and used without further purification.

### Preparation of ethanolic and aqueous propolis extracts (EEP and WEP)

Fresh propolis samples were collected from beehives in Chiang Mai Province, Thailand, in February 2019. Approximately 40 g of propolis was cut into small pieces, frozen in liquid nitrogen, and ground using a mortar and pestle. For the ethanolic extract (EEP), the ground propolis was soaked in 70% (v/v) ethanol for 24 h at room temperature with occasional stirring. For the aqueous extract (WEP), the same procedure was performed using distilled water as the solvent. The mixtures were filtered through Whatman No. 1 filter paper, and the filtrates were concentrated under reduced pressure using a rotary evaporator at 40 °C. The resulting dried extracts were weighed and stored at 4 °C until further analysis.

### DPPH free radical scavenging assay of EEP and WEP

The antioxidant activity of the EEP and WEP was determined using the 2,2-diphenyl-1-picrylhydrazyl (DPPH) radical scavenging assay, with minor modifications based on the method described by Isla et al.[17]. Extract solutions were prepared in ethanol at concentrations ranging from 0.5 to 5 mg/mL. The reaction mixture consisted of 2.4 mL of ethanol, 0.3 mL of 200 µM DPPH solution, and 0.3 mL of extract solution. The mixtures were vortexed and incubated at room temperature for 30 min in the dark. Absorbance was measured at 517 nm using a UV–Vis spectrophotometer against a control containing all reagents except the extract. All experiments were performed in triplicate. The percentage of DPPH radical scavenging activity was calculated using the following equation:

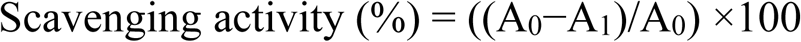

where A_0_ is the absorbance of the control (DPPH solution without extract), and A_1_ is the absorbance of the sample.

### Hydrogen peroxide scavenging assay of EEP and WEP

The hydrogen peroxide (H_2_O_2_) scavenging activity of the EEP and WEP was determined according to a modified spectrophotometric method [18]. Extract solutions were prepared in ethanol at concentrations ranging from 0.5 to 5 mg/mL. For each reaction, 2.4 mL of extract solution was mixed with 0.6 mL of H_2_O_2_ prepared in 0.1 M phosphate buffer (pH 7.4). The mixture was incubated at room temperature for 10 min, and the absorbance was measured at 230 nm using a UV–Vis spectrophotometer. Phosphate buffer without H_2_O_2_ served as the blank, and a solution without extract served as the control. The percentage of hydrogen peroxide scavenging activity was calculated using the equation:

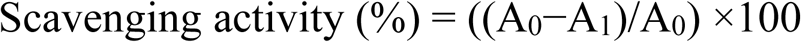

where A_0_ is the absorbance of the control and A_1_ is the absorbance of the sample. All experiments were performed in triplicate.

### Chemical profiling of EEP using gas chromatography–mass spectrometry (GC–MS)

The chemical composition of the ethanolic extract of propolis (EEP) was analyzed using gas chromatography–mass spectrometry (GC–MS) after derivatization to convert non-volatile compounds into volatile derivatives. The dried extract was mixed with N, O-bis(trimethylsilyl)trifluoroacetamide (BSTFA) containing 1% trimethylchlorosilane (TMCS) and incubated at 70 °C for 30 min to complete silylation. The derivatized sample was then analyzed using an Agilent 7890B GC–MS system equipped with an HP-5MS capillary column (30 m × 0.25 mm i.d., 0.25 µm film thickness). Helium was used as the carrier gas at a constant flow rate of 0.7 mL/min. The oven temperature was programmed from 100 °C to 300 °C at a rate of 5 °C/min and maintained for 5 min, with a total run time of 45 min. The injector temperature was set to 280 °C, and a 1 µL aliquot of the sample was injected in split mode (20:1). The mass spectrometer was operated in electron impact (EI) ionization mode at 70 eV. Compounds were identified by comparing their mass spectra with those in the Wiley 138 and NIST 98 spectral libraries.

### GC–MS data analysis and compound identification

The chromatographic and mass spectral data were processed using the Agilent MassHunter Workstation software. Each peak was identified based on a comparison of its mass spectrum with those available in the Wiley 138 and NIST 98 spectral libraries, considering only matches with a similarity index above 85%. The retention indices of selected compounds were compared with published data to support identification. Quantification was performed using the relative percentage of each compound, calculated from the ratio of its individual peak area to the total peak area of all detected components in the chromatogram. Identified compounds were further classified into major chemical groups, including flavonoids, phenolic acids, esters, and terpenes, based on their structural characteristics and literature reports on propolis chemistry.

### Hydroxyl radical scavenging assay of CAPE

The hydroxyl radical scavenging activity of CAPE was evaluated using the deoxyribose degradation method [19]. The reaction mixture contained 0.1 M sodium phosphate buffer (pH 7.0), 10 mM deoxyribose, 10 mM FeSO_4_–EDTA, and 1 mM H_2_O_2_. CAPE at various concentrations (2− 10 μM) was added to the mixture, and the reaction was incubated at 37 °C for 3 h. Following incubation, trichloroacetic acid and 2-thiobarbituric acid were added, and the mixture was boiled for 10 min to develop color. The absorbance was measured at 520 nm. The hydroxyl radical scavenging activity was calculated using the following equation:

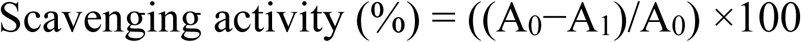

where A_0_ is the absorbance of the control (without CAPE) and A_1_ is the absorbance in the presence of CAPE. All experiments were performed in triplicate.

### Superoxide anion scavenging assay of caffeic acid phenethyl ester (CAPE)

The superoxide anion scavenging activity of CAPE was evaluated using a xanthine/xanthine oxidase (XOD) system [20]. The reaction mixture contained 0.1 mL of 3 mM xanthine, 0.1 mL of 3 mM EDTA, 0.1 mL of 0.15% (w/v) bovine serum albumin, and 0.1 mL of 0.75 mM nitroblue tetrazolium (NBT) in 0.05 M sodium carbonate buffer (pH 10). CAPE at various concentrations (2− 10 μM) was added to the mixture and incubated at room temperature for 10 min. The reaction was initiated by adding xanthine oxidase and allowed to proceed for 20 min. The reaction was terminated by the addition of 1 mL of 6 mM CuCl_2_, and the absorbance was measured at 560 nm. The percentage of superoxide anion scavenging activity was calculated using the following equation:

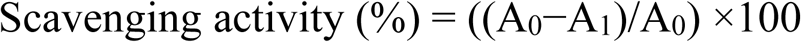

where A_0_ is the absorbance of the control (without CAPE) and A_1_ is the absorbance of the reaction mixture containing CAPE. All measurements were performed in triplicate.

### Protective effect of CAPE against plasmid DNA damage

The protective effect of CAPE against hydroxyl radical–induced DNA damage was evaluated using plasmid DNA (pBR322) [21]. The reaction mixture (total volume 25 µL) contained 1 M phosphate buffer (pH 7.4), 1 µg of pBR322 DNA, 100 µM CuCl₂, and 1 mM H_2_O_2_. CAPE at concentrations of 1, 2, and 4 µM was added to the reaction mixture to evaluate its protective effect on plasmid DNA. Quercetin (1 mM) served as the reference compound. After incubation for 30 minutes, the samples were analyzed by agarose gel electrophoresis at 100 V. DNA bands were visualized using ethidium bromide staining and compared with the control to evaluate the extent of DNA protection.

### Statistical Analysis

All experiments were performed in triplicate, and results are expressed as mean ± standard deviation (SD). Statistical comparisons were performed using one-way analysis of variance (ANOVA) and t-test in GraphPad Prism 10. Differences were considered statistically significant at *p* < 0.05.

## Results

### DPPH radical scavenging of EEP and WEP

The antioxidant activities of the EEP and WEP were assessed using the DPPH radical scavenging assay. EEP showed significantly higher inhibition than WEP at most concentrations (*p* < 0.05) (Fig. 1), whereas no significant difference was observed at the lowest concentration tested (0.5 mg/mL). The highest activities were observed at 4 and 5 mg/mL EEP, with inhibition percentages of 87.14% (t-test,p= 0.072) and 87.44% (t-test,p=0.055). WEP demonstrated moderate radical scavenging activity, ranging from 10.37-51.60 % of inhibition across the tested concentrations. Ascorbic acid (1 mM), used as a positive control, exhibited 81.95% inhibition.

**Figure 1.**
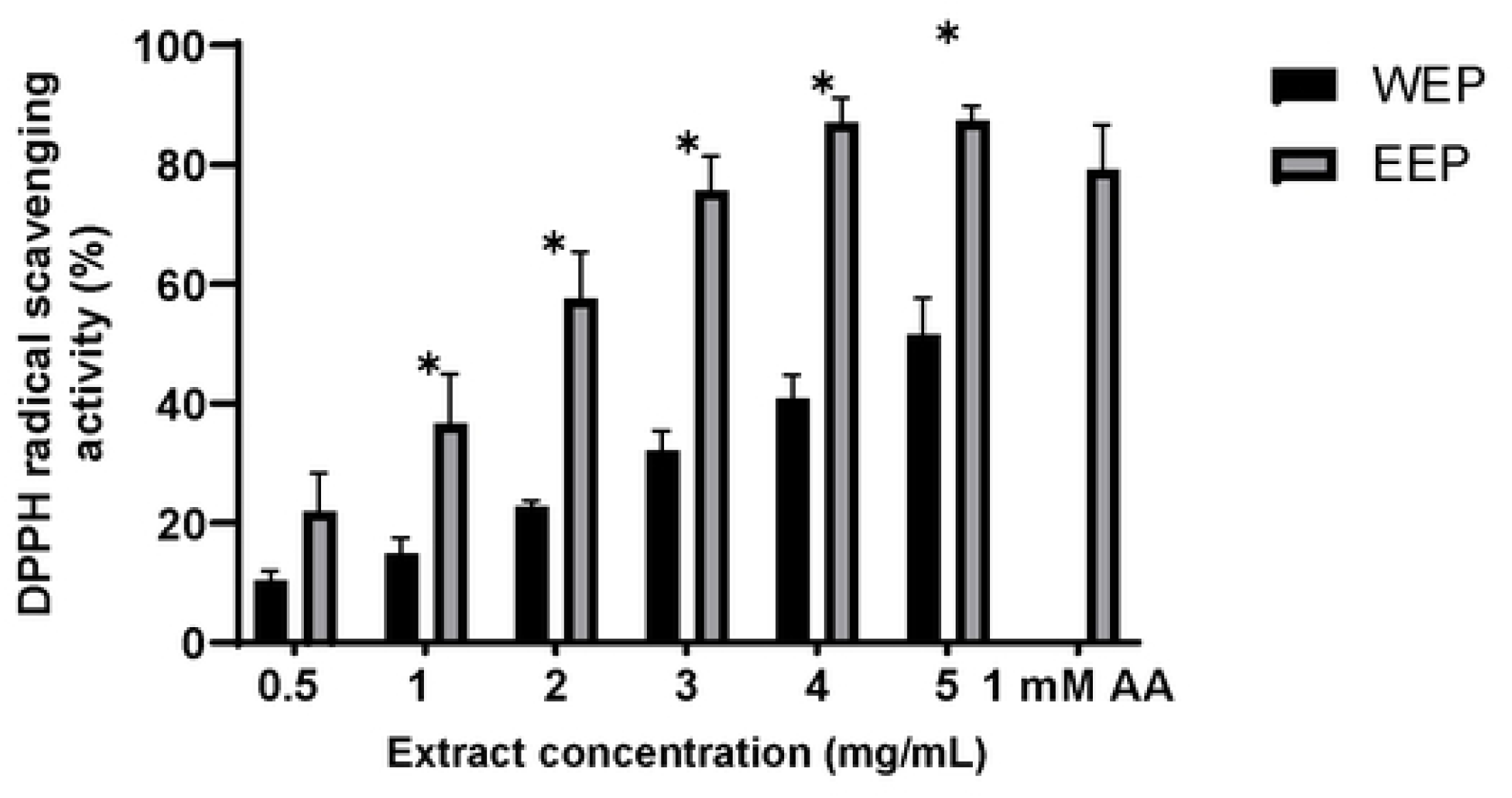
DPPH free radical scavenging of EEP and WEP. Data are represented mean and standard deviations (SD). P-value was significant at p < 0.05.

### Hydrogen peroxide scavenging activity of EEP and WEP

The hydrogen peroxide (H_2_O_2_) scavenging activity of EEP and WEP was assessed. Both extracts reduced H_2_O_2_ in a concentration-dependent manner (Fig. 2). No significant difference was observed between EEP and WEP in their overall H_2_O_2_ scavenging ability (*p* > 0.05). At the lower concentrations (0.5–1 mg/mL), WEP exhibited slightly higher scavenging activity than EEP, although the difference was not statistically significant. The scavenging activity of EEP ranged from 26.79 to 90.13%, with the highest activity observed at 4 mg/mL (90.13% inhibition). WEP showed inhibition ranging from 28.66 to 94.01% across the tested concentrations. EEP and WEP at concentrations of 4–5 mg/mL exhibited higher scavenging activity than 1 mM ascorbic acid, with inhibition values of 78.07%.

**Figure 2.**
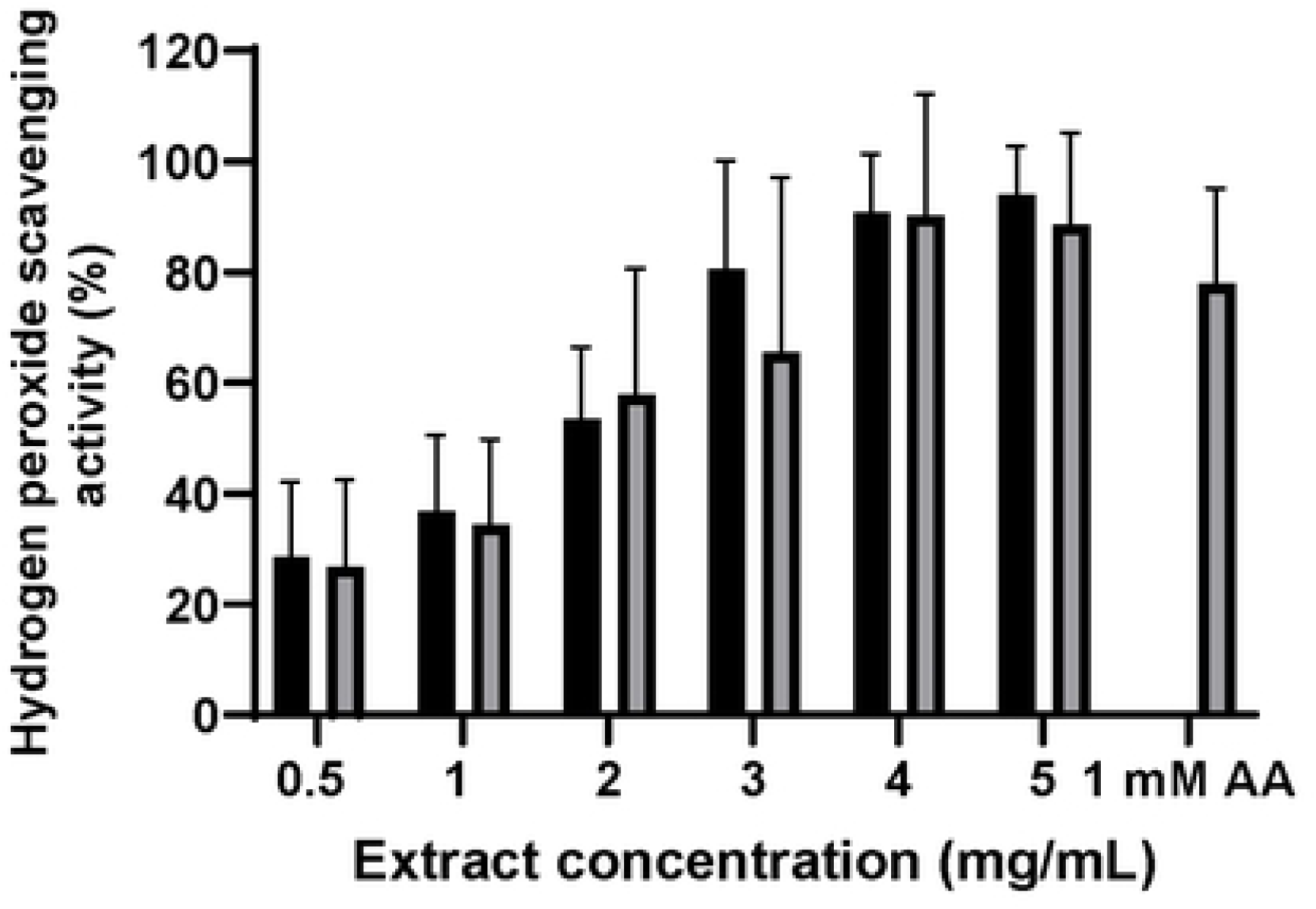
The inhibition of hydrogen peroxide of EEP and WEP. Data are represented mean and standard deviations (SD). P-value was significant at p < 0.05.

### Chemical profiling of EEP

The chemical constituents of the ethanolic extract of Thai propolis (EEP) were identified by gas chromatography–mass spectrometry (GC–MS) (Table 1 and Fig. 3). A total of 30 compounds were detected, comprising mainly phenolic acids, flavonoid derivatives, sugars, and their trimethylsilyl (TMS) derivatives. The most abundant constituents were D-altrose (13.17%), 2-hydroxybenzoic acid (6.48%), D-(−)-fructofuranose pentakis(trimethylsilyl) ether (6.24%), and galactonic acid, gamma-lactone (5.60%). Other notable compounds included 9,12-octadecadienoic acid (5.47%), L-altrose, 2,3,4,5,6-pentakis-O-(trimethylsilyl) (4.61%), and D-fructose (4.18%). Minor components identified were caffeic acid (0.13%), caffeic acid phenethyl ester (CAPE, 0.35%), p-coumaric acid (0.81%), E-cinnamyl Z-cinnamic (0.34%), and 3,4-dihydroxybenzoic acid (0.48%), which are known to contribute to the antioxidant and antimicrobial properties of propolis.

**Fig. 3.**
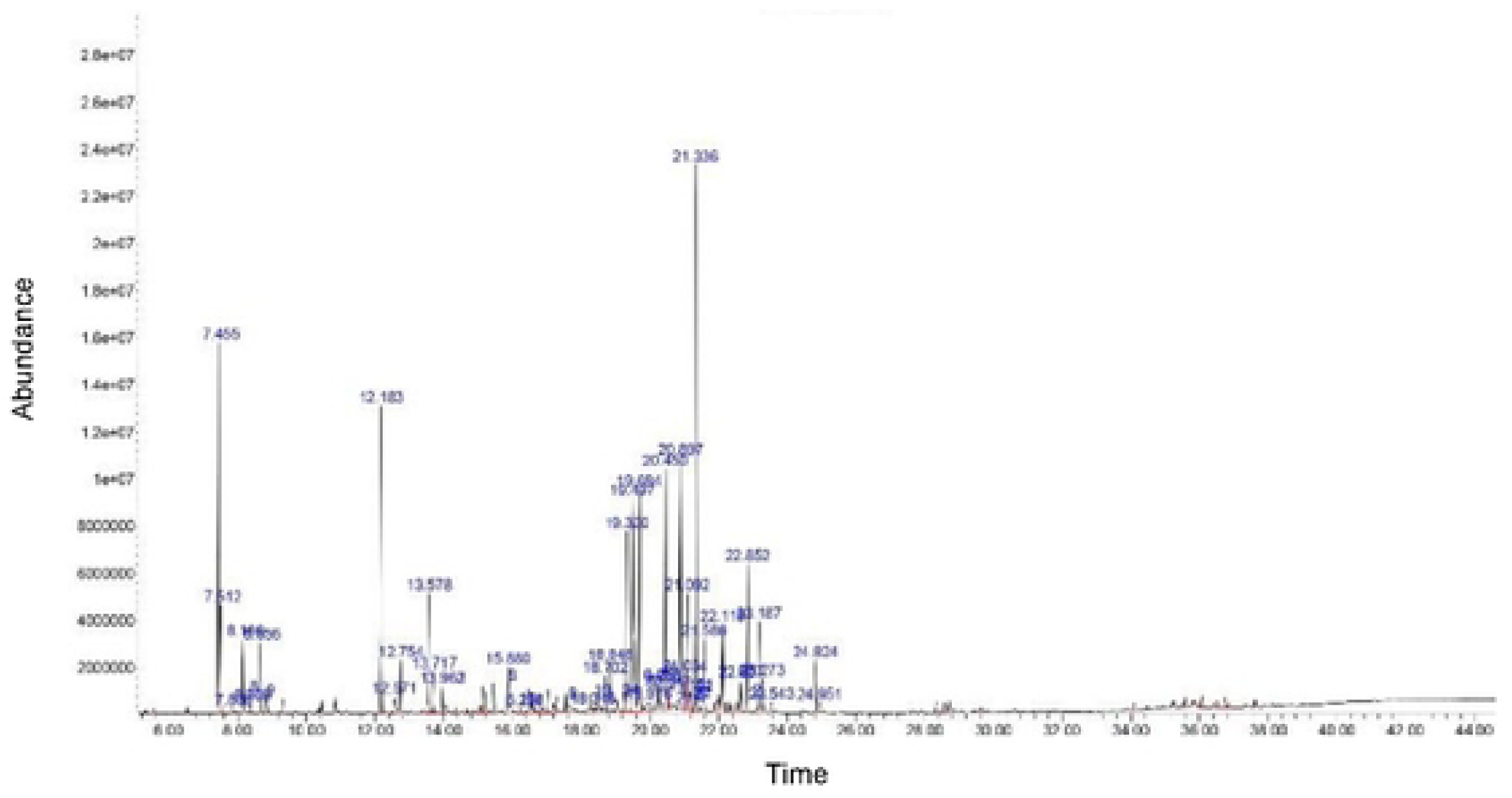
GC-MS analysis of EEP sample

**Table 1.**
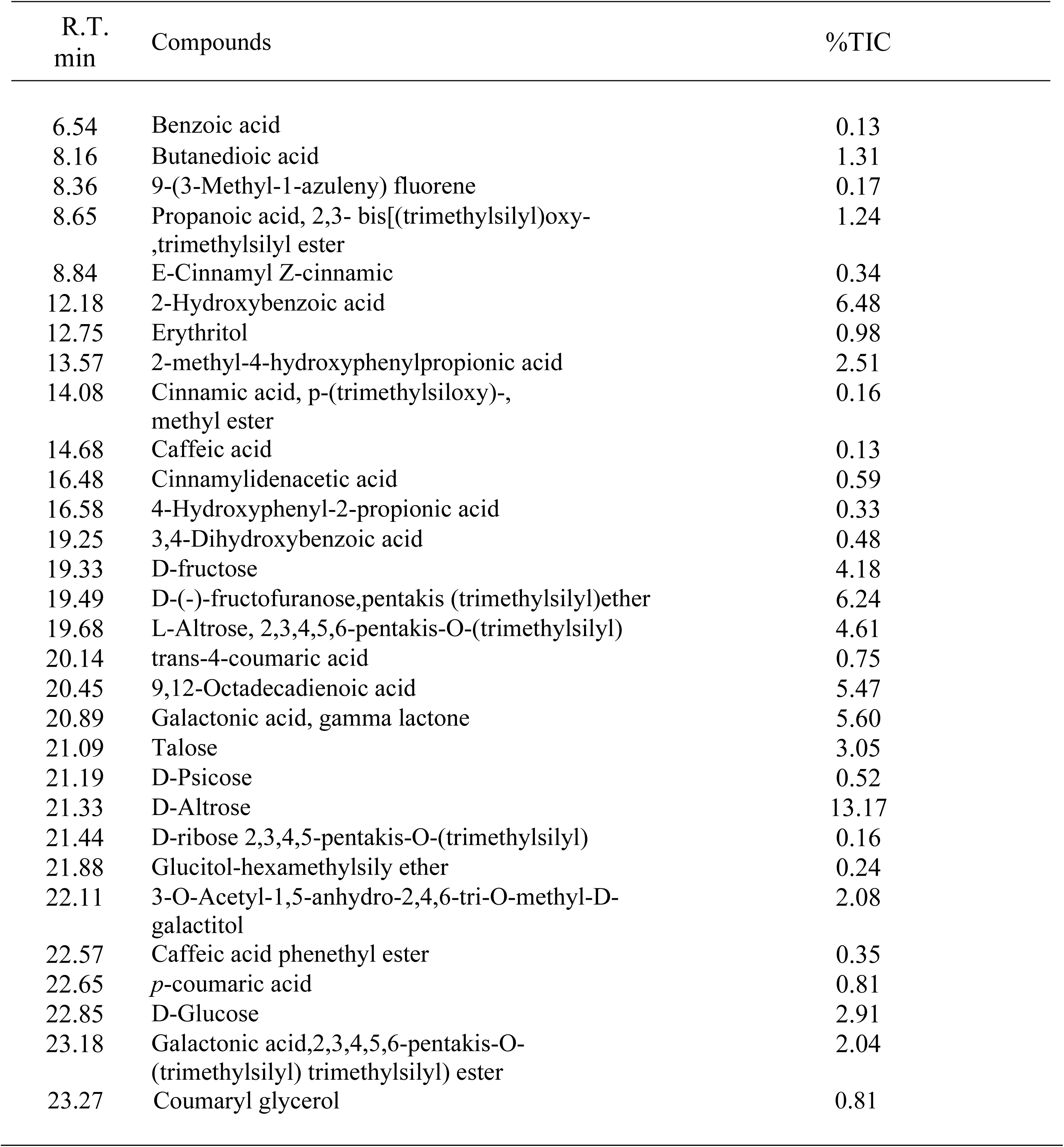
Compounds identified in propolis extract of EEP by GC-MS.

### Hydroxyl radical scavenging activity of CAPE

The hydroxyl radical scavenging activity of CAPE was evaluated, showing modest inhibition across the tested concentrations (2–10 µg/mL), with scavenging activity ranging from 16.54 to 57.56% (Fig. 4). In comparison, alpha-tocopherol exhibited strong hydroxyl radical inhibition, achieving 51.21% under the same conditions.

**Figure 4.**
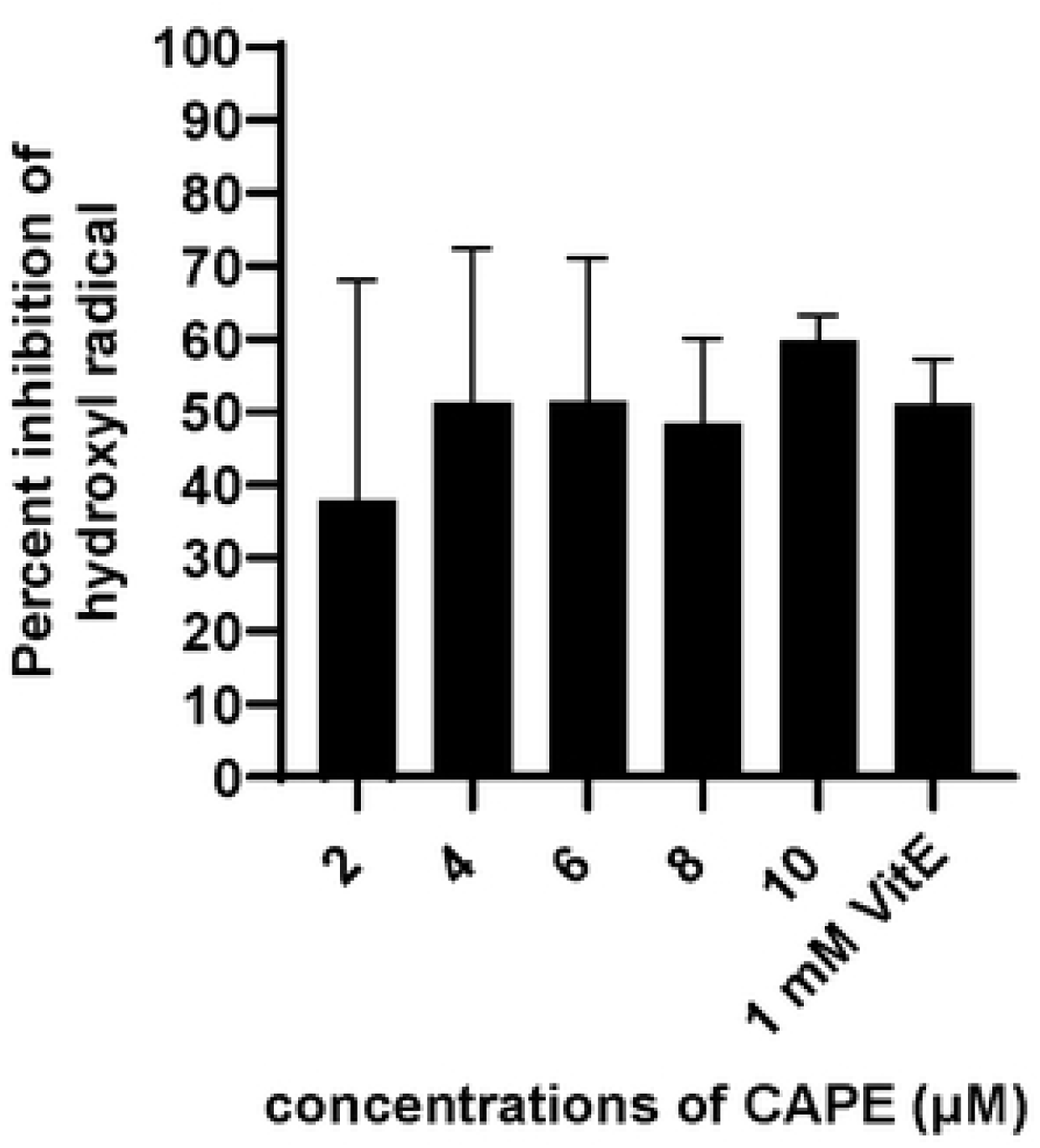
The inhibition of hydroxyl radical of CAPE. Data are represented mean and standard deviations (SD). P-value was significant at p ≤ 0.05.

### Superoxide anion scavenging activity of CAPE

The superoxide anion scavenging activity of caffeic acid phenethyl ester (CAPE) was assessed using the xanthine–xanthine oxidase system. CAPE inhibited superoxide anion generation in a dose-dependent manner, with higher concentrations exhibiting greater activity. At 10 µM, CAPE showed the highest scavenging activity, achieving 46.14% inhibition (Fig. 5). Lower concentrations of CAPE (2, 4, 6, and 8 µM) demonstrated scavenging activities of 3.29, 20.23, 21.88, and 35.63%, respectively.

**Figure 5.**
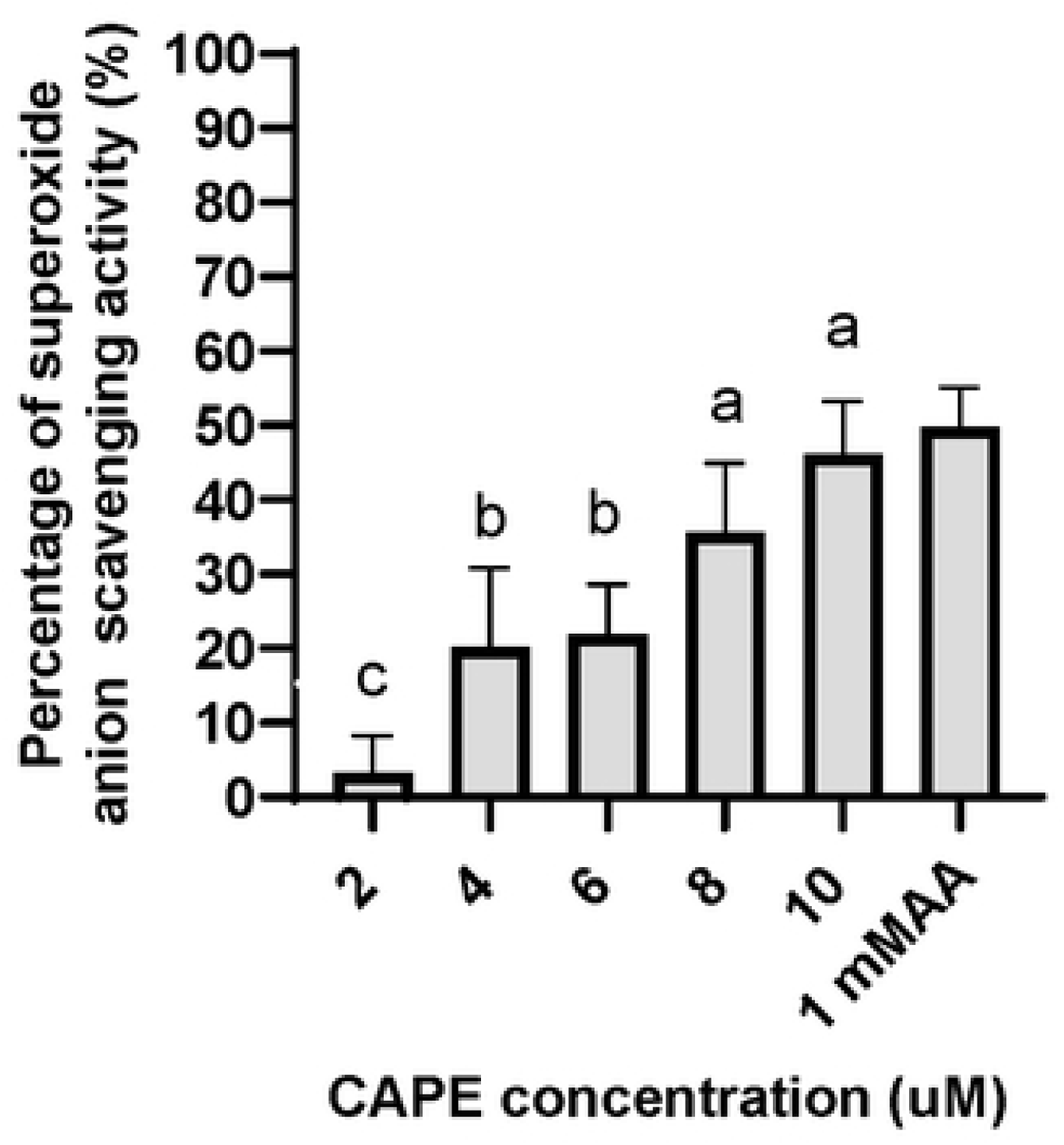
The inhibition of superoxide anion of CAPE. Data are represented mean and standard deviations (SD). P-value was significant at p ≤ 0.05.

### Plasmid DNA Protection by CAPE

The protective effect of CAPE on plasmid DNA damage was evaluated using the pBR322 plasmid system (Fig. 6). DNA strand breakage was induced by hydrogen peroxide (H_2_O_2_) in the presence of copper ions, and the ability of CAPE to prevent oxidative degradation was assessed. In the control lane (Lane 1), plasmid DNA treated with 1 µM CAPE alone maintained an intact supercoiled form, indicating no DNA degradation. Upon exposure to H_2_O_2_ (Lanes 2 and 3), the presence of CAPE at 2 and 4 µM reduced the conversion of supercoiled DNA to open circular or linear forms. In contrast, quercetin (1 mM) with H_2_O_2_ (Lane 4) exhibited greater DNA nicking and linearization, indicating less effective protection under these conditions.

**Figure 6.**
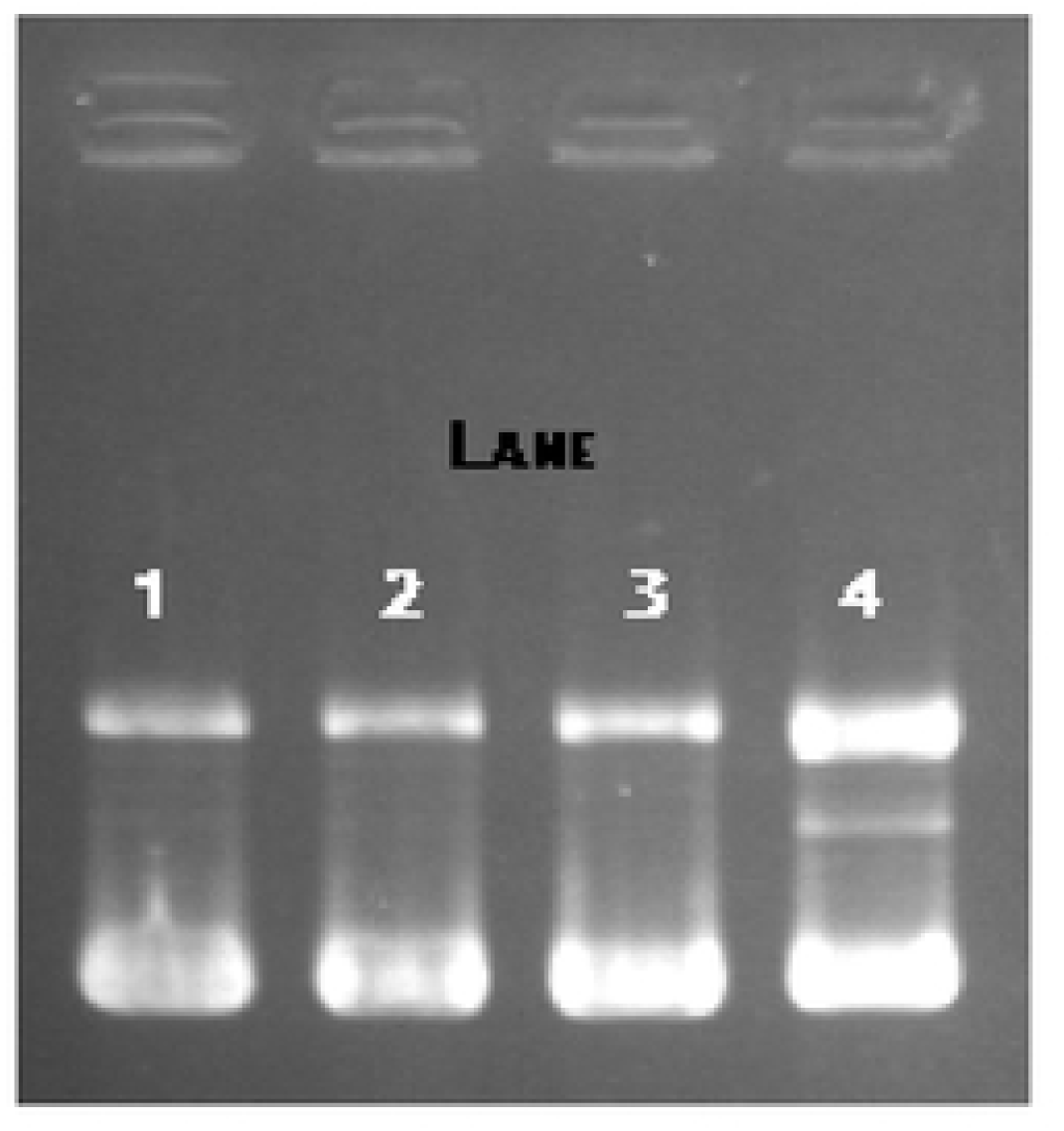
Protection of DNA breakage with CAPE. Plasmid DNA with CAPE in 1 % agarose gel staining with 1 mg/ml EtBr; Lane 1 pBR322 1 µg + 1 µM CAPE + PBS 10 µl +100 µM CuCl_2_ Lane 2 pBR322 1 µg + 2 µM CAPE + PBS 10 µl +100 µM CuCl_2_ + 1 mM H_2_O_2_ Lane 3 pBR322 1 µg + 4 µM CAPE + PBS 10 µl +100 µM CuCl_2_ + 1 mM H_2_O_2_ Lane 4 pBR322 1 µg + 1 mM QT + PBS 10 µl +100 µM CuCl_2_ + 1 mM H_2_O_2_

## Discussion

In this study, both ethanolic (EEP) and aqueous (WEP) extracts of Thai propolis exhibited significant antioxidant activity, confirming the presence of phenolic and flavonoid compounds as the main bioactive constituents. EEP demonstrated higher radical scavenging activity than WEP, consistent with the solubility of phenolic compounds in ethanol. This finding supports previous research showing that ethanolic extraction yields a broader spectrum of antioxidant compounds from propolis compared to aqueous extraction [22, 23,24]. The stronger inhibition of DPPH radicals and hydrogen peroxide observed in EEP highlights its potential as an effective natural antioxidant source.

Hydrogen peroxide and superoxide anions are reactive oxygen species (ROS) that can cause oxidative stress, leading to lipid peroxidation, DNA damage, and cellular dysfunction. Both EEP and WEP effectively scavenged hydrogen peroxide, indicating that Thai propolis contains both polar and non-polar antioxidant compounds capable of neutralizing ROS [25]. Although WEP displayed lower activity overall, its antioxidant potential at higher concentrations suggests that some hydrophilic phenolics are also responsible for the activity in the aqueous fraction. These results demonstrate that both types of extracts could contribute to cellular protection against oxidative damage. Caffeic acid phenethyl ester (CAPE), identified as one of the active compounds in Thai EEP, exhibited strong antioxidant potential through its superoxide and hydroxyl radical scavenging abilities. CAPE efficiently inhibited superoxide generation in the xanthine–xanthine oxidase system and reduced hydroxyl radicals, confirming its dual role as a free radical scavenger and metal chelator. Similar findings have been reported in CAPE isolated from other propolis sources, where it showed protective effects against H₂O₂-induced oxidative stress in neuronal and epithelial cells and improved cell viability by modulating redox balance [26]. The plasmid DNA protection assay further revealed that CAPE mitigated oxidative DNA damage induced by Cu²⁺ and H₂O₂, supporting its potential as a cytoprotective compound. In contrast, quercetin, used as a reference flavonoid, promoted plasmid breakage at high concentrations, suggesting possible prooxidant effects under certain conditions.

The chemical composition of Thai propolis observed in this study shares similarities with propolis from other geographical origins but also shows unique features. For example, Brazilian green propolis is rich in prenylated p-coumaric acid derivatives, artepillin C, and cinnamic acid derivatives [27, 28], while Taiwanese propolis mainly contains prenylated flavanones [29], and Cuban propolis possesses unusual polyprenylated benzophenones with cytotoxic effects on cancer cells [30]. Portuguese propolis, on the other hand, contains phenolic compounds associated with anti-melanoma and antitumor activities [31]. In the present study, Thai propolis contained caffeic acid, cinnamic acid derivatives, flavanones, sugars, and prenylated phenolic compounds, including caffeic acid phenethyl ester (CAPE), confirming its complex and diverse phytochemical profile. These compounds are known for their antioxidant, antimicrobial, and cytoprotective properties, suggesting that Thai propolis shares the functional potential of propolis from other regions while possessing distinct compositional characteristics influenced by local flora.

The strong antioxidant activities of EEP and CAPE indicate that Thai propolis could serve as a promising source of natural antioxidants and bioactive molecules for pharmaceutical, nutraceutical, and food applications. Its efficacy in scavenging free radicals and protecting DNA against oxidative damage highlights its potential role in preventing oxidative stress–related diseases. Further research is warranted to explore the detailed mechanisms of CAPE and other active constituents in Thai propolis, as well as their synergistic effects and stability in biological systems.

## Conclusion

In conclusion, Thai propolis contains diverse bioactive compounds, particularly polyphenols and flavonoids, that contribute to its strong antioxidant capacity. The ethanolic extract (EEP) demonstrated superior radical scavenging activity compared to the aqueous extract (WEP), primarily due to the presence of caffeic acid phenethyl ester (CAPE) and related phenolic derivatives. CAPE effectively scavenged superoxide and hydroxyl radicals and protected plasmid DNA from oxidative damage, highlighting its potential as a natural antioxidant and cytoprotective agent. These findings suggest that Thai propolis may serve as a valuable source of functional compounds for applications in food preservation, nutraceutical development, and oxidative stress–related disease prevention.

## Data availability

The datasets used and/or analyzed during the current study are available from the corresponding author on reasonable request.

## Acknowledgements

This research was financially supported by Thailand Science Research and Innovation and Maejo University, Thailand (grant number MJU 1-61-117).

## Author Contributions

P.P. conceived and designed the experiments, and conducted the research. T.C. performed the GC–MS analysis. P.P. and T.C. analyzed the data and wrote the manuscript. J.J. and V.C. reviewed and edited the manuscript. All authors have read and approved the final version of the manuscript.

## Additional information

The corresponding author is responsible for submitting a competing interests statement on behalf of all authors of the paper.

## References

1. Bankova V, Popova M, Trusheva B. The phytochemistry of the honeybee. Phytochemistry. 2018; 155:1–11. 10.1016/j.phytochem.2018.07.007.

2. Salatino A, Teixeira EW, Negri G, Message D. Origin and chemical variation of propolis. eCAM. 2005; 2 (1): 33–38. 10.1093/ecam/neh060.

3. Kosalec I, Bakmaz M, Pepeljnjak S, Ladimir-Knezević SV. Quantitative analysis of the flavonoids in raw propolis from northern Croatia. Acta Pharm. 2004; 54:65–72.

4. Miłek M, Ciszkowicz E, Tomczyk M, Sidor E, Zaguła G, Lecka-Szlachta K, Pasternakiewicz A, Dżugan M. The study of chemical profile and antioxidant properties of poplar-type polish propolis considering local flora diversity in relation to antibacterial and anticancer activities in human breast cancer cells. Molecules. 2022; 27 (3) :725–747. 10.3390/molecules27030725.

5. Velikova M, Bankova V, Marcucci MC, Tsvetkova I, Kujumgiev A. Chemical composition and biological activity of propolis from Brazilian Meliponinae. Z Naturforsch C J Biosci. 2000;55 (9-10): 785–789. 10.1515/znc-2000-9-1018.

6. Kumazawa S, Yoneda M, Shibata I, Kanaeda J, Hamasaka T, Nakayama T. Direct evidence for the plant origin of Brazilian propolis by the observation of honeybee behavior and phytochemical analysis. Chem Pharm Bull Tokyo. 2003; 51(6):740–742. 10.1248/cpb.51.740.

7. Kim HG, Han EH, Im JH, Lee EJ, Jin SW, Jeong HG. Caffeic acid phenethyl ester inhibits 3-MC-induced CYP1A1 expression through induction of hypoxia-inducible factor-1α. Biochem Biophys Res Commun. 2015; 465 (3): 562–568. 10.1016/j.bbrc.2015.08.060.

8. Derman S. Caffeic acid phenethyl ester loaded PLGA nanoparticles: Effect of various process parameters on reaction yield, encapsulation efficiency, and particle size. J Nanomater. 2015; 341848: 1–15. 10.1155/2015/341848.

9. Tseng J, Lin C, Su L, Fu H, Yang S, Chuu C. CAPE suppresses migration and invasion of prostate cancer cells via activation of non-canonical Wnt signaling. Oncotarget. 2016; 7: 38010–38024. 10.18632/oncotarget.9380.

10. Banskota AH, Nagaoka T, Sumioka LY, Tezuka Y, Awale S, Midorikawa K, Matsushige K, Kadota S. Antiproliferative activity of the Netherlands propolis and its active principles in cancer cell lines. J Ethnopharmacol. 2002; 80:67–73. 10.1016/s0378-8741(02)00022-3.

11. Kantrong N, Kumtawee J, Damrongrungruang T, Puasiri S, Makeudom A, Krisanaprakornkit S, Chailertvanitkul P. An in vitro anti-inflammatory effect of Thai propolis in human dental pulp cells. J Appl Oral Sci. 2023; 31:e20230006. 10.1590/1678-7757-2023-0006.

12. Tatlısulu S, Özgör E. Identification of Cyprus propolis composition and evaluation of its antimicrobial and antiproliferative activities. Food Biosci. 2023;51:102273–102281. 10.1016/j.fbio.2022.102273.

13. Moreno MA, Vallejo AM, Ballester AR, Zampini C, Isla MI, López-Rubio A, Fabra MJ. Antifungal edible coatings containing Argentinian propolis extract and their application in raspberries. Food Hydrocoll. 2020;107: 105973–105982. 10.1016/j.foodhyd.2020.105973.

14. Trusheva B, Trunkova D, Bankova V. Different extraction methods of biologically active components from propolis: a preliminary study. Chem Cent J. 2007; 1:1–13. 10.1186/1752-153X-1-13.

15. Zeng L, Xia T, Hu W, Chen S, Chi S, Lei Y, Liu Z. Visualizing the regulation of hydroxyl radical level by superoxide dismutase via a specific molecular probe. Anal Chem. 2018;90(2) :1317–1324. 10.1021/acs.analchem.7b04191.

16. Callahan LA, She ZW, Nosek TM. Superoxide, hydroxyl radical, and hydrogen peroxide effects on single-diaphragm fiber contractile apparatus. J Appl Physiol. 2001; 90(1):45–54. 10.1152/jappl.2001.90.1.45.

17. Isla MI, Nieva Moreno MI, Sampietro AR, Vattuone MA. Antioxidant activity of Argentine propolis extracts. J Ethnopharmacol. 2001;76(2):165–70. doi: 10.1016/s0378-8741(01)00231-8.

18. Ak T, Gülçin I. Antioxidant and radical scavenging properties of curcumin. Chem Biol Interact. 2008;174(1):27–37. doi: 10.1016/j.cbi.2008.05.003.

19. Joubert E, Winterton P, Britz TJ, Gelderblom WC. Antioxidant and pro-oxidant activities of aqueous extracts and crude polyphenolic fractions of rooibos (Aspalathus linearis). J Agric Food Chem. 2005;53(26):10260–7. doi: 10.1021/jf051355a.

20. Pascual C, Gonzalez R, Torricella RG. Scavenging action of propolis extract against oxygen radicals. J Ethnopharmacol. 1994;41(1-2):9–13. doi: 10.1016/0378-8741(94)90052-3.

21. Guleria, S., Tiku, A.K., Singh, G., Vyas, D. and Bhardwaj, A. (2011), Antioxidant Activity and Protective Effect Against Plasmid DNA Strand Scission of Leaf, Bark, and Heartwood Extracts from *Acacia catechu*. Journal of Food Science, 76: C959–C964. 10.1111/j.1750-3841.2011.02284.x

22. Ahn M, Kumazawa S, Usui Y, Nakamura J, Matsuka M, Zhu F, Nakayama T. Antioxidant activity and constituents of propolis collected in various areas of China. Food Chem. 2007;101 (4): 1383–1392. 10.1016/j.foodchem.2006.03.045.

23. Nagai T, Inoue R, Inuoe H, Suzuki N. Preparation and antioxidant properties of water extract of propolis. Food Chem. 2003; 80 (4): 29–33. 10.1016/S0308-8146(02)00231-5.

24. Sevgi K, Ceren B. A comparative study of solvent effect on propolis extraction by ultrasound assisted extraction, Turk J Anal Chem. 2024; 6 (1): 11–17. 10.51435/turkjac.1445121.

25. Pukklay P, Chuesaard T. Total phenolic, flavonoid contents and antioxidant activity of propolis extracts from Nan province. PSRU J Sci Technol 2021;6:13–27.

26. Lai X, Najafi M. Redox interactions in chemo/radiation therapy-induced lung toxicity; mechanisms and therapy perspectives. Curr Drug Targets. 2022;23 (13):1261–1276. doi: 10.2174/1389450123666220705123315.

27. Park YK, Alencar SM, Aguiar CL. Botanical origin and Chemical composition of Brazilian propolis, J Agric Food Chem. 2002; 50 (9) :2502–2506. 10.1021/jf011432b.

28. Marcucci MC, Bankova V. Chemical Composition, Plant Origin and Biological Activity of Brazilian Propolis, Curr Top Phytochem. 1999; 2 ;115–123.

29. Chen CN, Wu CL, Shy HS, Lin JK. Cytotoxic prenylflavanones from Taiwanese propolis, J Nat Prod. 2003; 66 (4): 503–506. doi: 10.1021/np0203180.

30. Cuestro-Rubio O, Frontana-Uribe BA, Ramírez-Apan T, Cárdenas J. Polyisoprenylated benzophenones in Cuban Propolis;Biological Activity of Nemorosone. Z. Naturforsch C J Biosci. 2002; 57 (3-4): 372–378. doi: 10.1515/znc-2002-3-429.

31. Caetano AR, Oliveira RD, Celeiro SP, Freitas AS, Cardoso SM, Goncalves MST, Baltazar F, Almeida-Aguiar C. Phenolic Compounds Contribution to Portuguese Propolis Anti-Melanoma Activity. Molecules. 2023; 28 (7): 3107–3120. 10.3390/molecules28073107.

